# Cellpanelr: identify predictive biomarkers from cell line panel response data

**DOI:** 10.1101/2022.11.02.514913

**Authors:** Douglas R. Wassarman, Taiasean Wu, Kevan M. Shokat

## Abstract

**Summary:** Cellpanelr is an open-source R package and web application for analyzing user-generated cell panel screens using DepMap data sets. Cellpanelr can be used to identify mutation and expression biomarkers of cell line response, increasing the value and accessibility of cell panel experiments such as relative sensitivities to cancer drugs.

**Availability and implementation:** Hosted web application is available from shinyapps.io (https://dwassarman.shinyapps.io/cellpanelr). Source code and installation instructions are available from GitHub (https://github.com/dwassarman/cellpanelr).

## Introduction

The Cancer Dependency Map (DepMap) is a database that curates data sets of gene expression, mutations, gene essentiality, and drug sensitivity for over 1,000 cell lines^1,2^. Integration of these data to analyze cell panel screens has enabled the discovery of potential therapeutic targets, new gene interactions, and drug resistance mechanisms^3–5^. However, there remains a high barrier of entry for many researchers who wish to analyze new cell panel experiments, such as drug response, from these cell lines using these pre-existing data sets. The new data must be cleaned and merged with the existing data, followed by many, often thousands, of calculations and analyses. These steps are tedious and error-prone to perform manually, necessitating a programmatic solution.

To meet this need, we developed cellpanelr, an online analysis tool and R Shiny application^6,7^ that allows users to identify predictive biomarkers of cell line response using data sets available from DepMap (Fig. 1). The online Shiny application provides an intuitive and convenient graphical user interface for researchers to upload cell response data, find predictive biomarkers, and download results and visualizations. The R package allows users to run the Shiny application locally and provides convenience functions for analysis and data access that can be used inside other R code.

**Figure 1.**
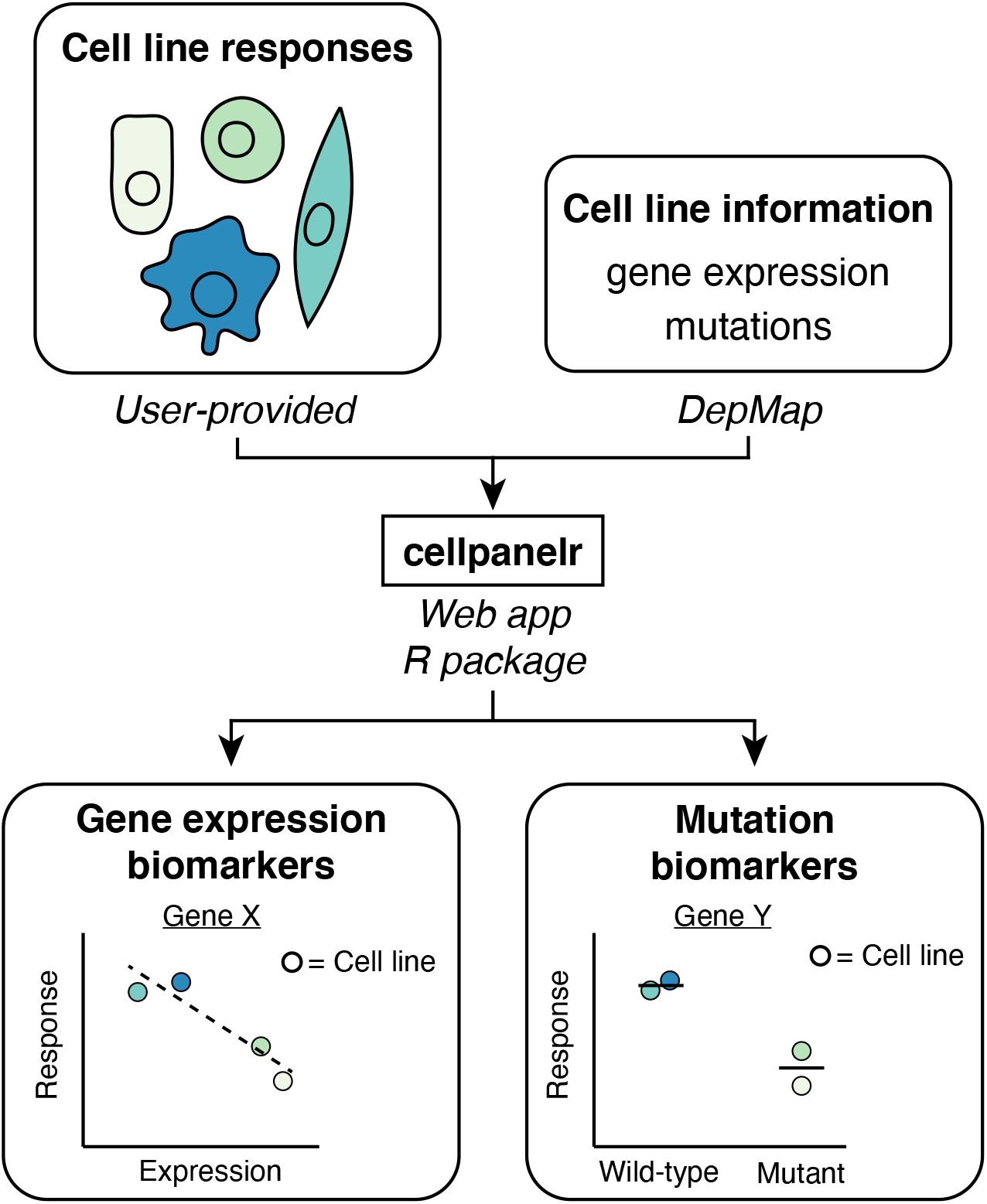
Cellpanelr analysis workflow. Users input response measurements from multiple cell lines using the interactive cellpanelr web app or R package. Expression or mutation biomarkers for the input response are determined using data sets available from DepMap.

## Methods

Cellpanelr was developed as an all-in-one Shiny/R package using the golem package and framework^8,9^. It makes use of tidyverse packages for data handling and visualizations^10^. Cell line annotations, gene expression, and mutation data sets were obtained from DepMap (https://depmap.org) and all analyses were performed using the 22Q1 release. Nultin-3 dose-response data was obtained from The Genomics of Drug Sensitivity in Cancer project (GDSC) (https://cancerrxgene.org)^11^.

User-provided cell line names are mapped to unique DepMap IDs using capitalized alpha-numeric characters. These IDs are used to merge user-provided response values with the gene expression, mutations, and annotations for each cell line. For each of 19,177 genes, Spearman’s correlation is calculated between cell line response and gene expression (log_2_[TPM + 1], TPM = transcripts per million) across cell lines. For each of 19,537 genes, Mann-Whitney U test is performed to determine whether cell lines with a mutation in the given gene respond differently than wild-type cell lines. Statistical significance for this analysis is determined using the Benjamini-Hochberg method with a false discovery rate of 0.05^12^.

## Results

To demonstrate and evaluate the cellpanelr workflow, we looked for biomarkers of cell line sensitivity to nutlin-3. Nutlin-3 is an inhibitor of the E3 ubiquitin ligase MDM2^13^. Nutlin-3 prevents MDM2-mediated degradation of the tumor suppressor p53, leading to an activation of the p53 pathway and cell cycle arrest. The nutlin-3 data set previously collected by GDSC contains IC_50_ and area under the dose-response curve (AUC) measurements for 968 cancer cell lines^11^. We elected to use AUC measurements for the cellpanelr analyses because many of the calculated IC_50_ values fall outside the tested dose range.

Of the cell lines in the nutlin-3 data set, gene expression data was available from DepMap for 680. We determined the Spearman correlation of each gene’s expression level with nutlin-3 sensitivity. Eleven of the top 12 genes most associated with nutlin-3 sensitivity are direct transcriptional targets of p53 or are known to be associated with p53 activation^14^ (Supp. Fig. S1). These genes include DDB2 (ρ = -0.42), BAX (ρ = -0.31), CCNG1 (ρ = -0.30), and MDM2 (ρ = - 0.37) itself. This indicates that cells with high p53 pathway activity are more sensitive to nutlin-3, consistent with the drug’s mechanism of action and its previously described selectivity profile^13^.

We then analyzed the AUC data set with cellpanelr to identify mutations that may be associated with nutlin-3 sensitivity or resistance. Mutation annotations were available from DepMap for 931 of the cell lines tested with nutlin-3. As expected, we found that mutations to p53 were highly associated with resistance to nutlin-3 (Supp. Fig. S2, p = 1.23 × 10^−65^). This result is consistent with existing knowledge that inhibition of p53 degradation does not reactivate the pathway in a mutant p53 context^13,15^. Furthermore, we confirmed prior studies that mutations to the tumor suppressor RB1 were associated with nutlin-3 resistance (p = 1.29 × 10^−8^, Supp. Fig. S2)^16^. Downstream activation of Rb is a critical aspect of p53-dependent cell cycle arrest. In Rb mutant or null cells, this activation cannot occur and the cells are therefore less responsive to nutlin-3.

In addition to recapitulating known resistance mutations to nutlin-3, cellpanelr identified a potential novel sensitizing mutation to nutlin-3. Cell lines harboring mutations to EH-domain containing protein 1 (EHD1) were significantly more sensitive to nutlin-3 (p = 6.43 × 10^−6^, Supp. Fig. S2). This result is interesting, as EHD1 has been previously described as a tumor suppressor in breast cancer which is involved with EGFR recycling^17^. To our knowledge, this is the first report linking EHD1 mutation with nutlin-3 sensitivity. More investigation is needed to validate and characterize the mechanism behind this observation.

## Discussion

Even with the availability of DepMap’s data sets, “-omics” level biomarker analyses remain inaccessible to many researchers. Cellpanelr lowers the barrier to these analyses through an interactive web application and R package. Users can upload response data from a collection of cell lines and, in minutes, have a list of genes that are associated with the given response along with interactive plots and visualizations. In this case study, we showed that cellpanelr can be easily used to recapitulate known gene expression and mutation biomarkers of drug sensitivity. Cellpanelr can accept any type of user-generated cell line response data, for example fluorescence, infection rate, or viability, and can therefore be applied to identify biomarkers of any quantitative cell line response. The case study on nutlin-3 sensitivity described here used input data containing hundreds of cell lines. Additional study is needed to determine the minimum number of cell lines needed to reliably detect biomarkers.

Cellpanelr implements several improvements to DepMap’s data portal. Cellpanelr does not require input cell lines to be pre-labeled with DepMap ID numbers, and instead performs this identification from standard cell line names. Input data can contain multiple response columns, allowing the user to select which response data to analyze. The DepMap portal uses Pearson’s correlation for determining relationships to gene expression and mutations, which is applicable only to linear correlations. Cellpanelr uses Spearman’s correlation for gene expression because it is robust to non-linear correlations. To analyze mutations, cellpanelr uses the Mann-Whitney U test instead of Pearson’s correlation because Mann-Whitney U is more appropriate for comparing two populations and, unlike the t-test, does not make assumptions about the shape of the population distributions. Cellpanelr is also open source and extensible, allowing users to access the underlying code to build upon cellpanelr’s functionality.

## Acknowledgements

This work was supported by the National Institutes of Health [5F31CA243439 to D.R.W]; and the Howard Hughes Medical Institute. Thank you to Megan Moore and Keelan Guiley for their feedback throughout the development of cellpanelr and preparation of this manuscript.

## Supplemental Figures

**Supplemental Figure S1.**
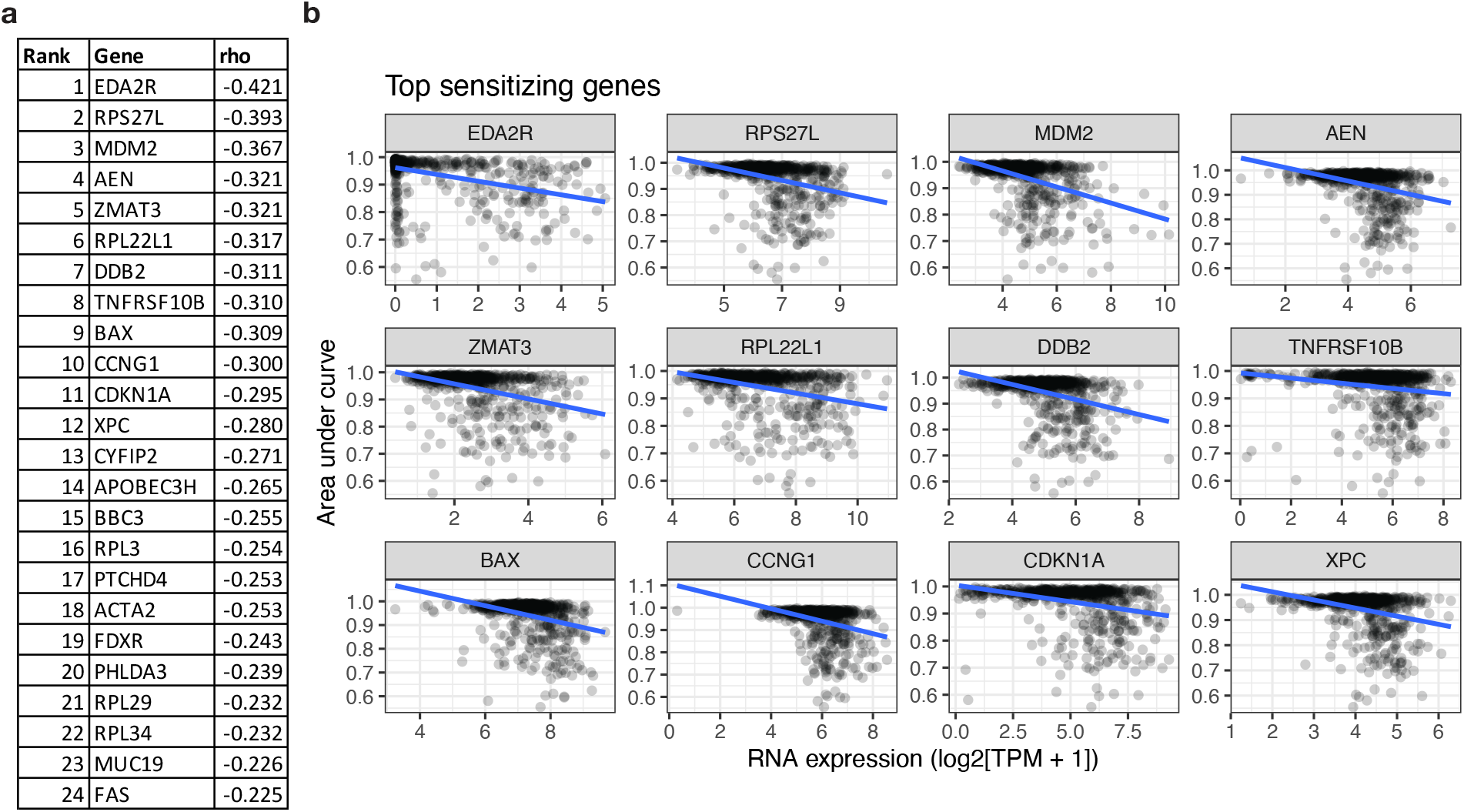
Gene expression associations with nutlin-3 sensitivity. (a) Top 24 genes associated with nutlin-3 sensitivity as determined by Spearman’s correlation (rho). (b) Gene expression (log-transformed transcripts-per-million) vs. area under the dose response curve for 12 genes with strongest association. Each dot represents a single cell line (n = 680). Blue line represents Pearson’s correlation.

**Supplemental Figure S2.**
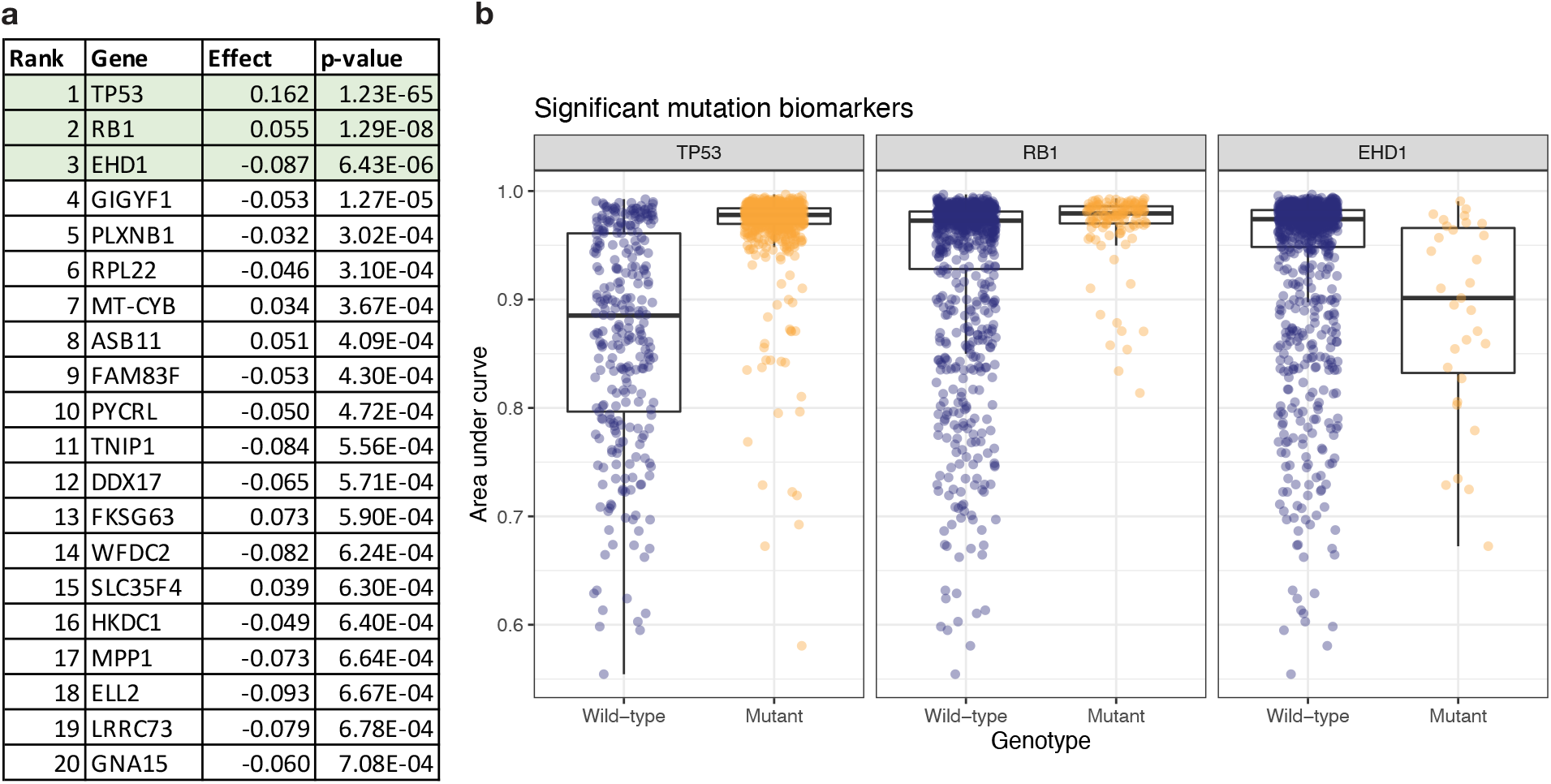
Mutations associated with nutlin-3 sensitivity and resistance. (a) Top 20 mutations most strongly associated with differences in nutlin-3 sensitivity as determined by Mann-Whitney U test. Statistically significant (false discovery rate ≤ 0.05) mutations highlighted in green. Effect calculated as log_2_(AUC_Mutant_ / AUC_Wild-type_). (b) Gene mutant status vs. area under the dose response curve for the three significant mutations. Each dot represents a single cell line (n = 931). Boxplot denotes the median and middle two quartiles, and whiskers show a maximum distance of 1.5 times this inner quartile range.

